# Novel Interpretable Tissue-Specific and Multi-Tissue Transcriptomic Clocks to Infer Aging Mechanisms

**DOI:** 10.1101/2021.05.11.443707

**Authors:** Aayush Gupta, Mindren Lu, Jessica Sun

## Abstract

Aging is characterized as a progressive decline in fitness that ultimately results in death. We set out to build both tissue-specific and multi-tissue transcriptomic clocks to make global tissue age predictions in individuals from GTEx. Existing work in the field primarily uses epigenetic clocks as predictors of age, but these models have known issues and are significantly less interpretable than their transcriptomic counterparts. Due to their transcriptomic nature, we can use these models to directly infer mechanisms of aging from their features. Linear regression remains the current standard analysis technique, but we improved upon its baseline performance with modern techniques, exploring both XGBoost and MLPs. We also experimented with using deconvolved cell data for predictions, which account for cellular composition and reduce signal distortion from rare cell types. Since it is known that the heterogeneity of cell types in particular tissues can lead to noise in these models, we proposed using deconvolution as a potential remedy for this problem.

Our results found that MLPs are not well suited for the task due to a lack of training data, but the use of XGBoost is effective at improving the baseline performance of predictions of existing tissue-specific clocks. These models allowed us to directly compute genes most important to age prediction in our models, and we showed that multiple genes found have been independently identified elsewhere to show evidence of correlation with age. Given the small size of our datasets, we were unable to make conclusive determinations about multi-tissue predictors, but preliminary results suggest that the technique shows promise and is worthy of future investigation. Likewise, given our limited deconvolved cell data, we did not currently observe strong results, but we again note that this is an area in need of further investigation.

By improving upon the performance of existing models, we demonstrated that a novel machine learning technique, XGBoost, can be an effective technique to further our understanding of aging mechanisms by extraction of the most relevant genes found in those models. This is significant because the genetic causes of aging are still not fully understood, and research in the field of aging is lacking in comparison to other domains. As the problem of identifying tissues that age at different rates is of specific interest, our tissue-specific models potentially have other applications in this domain, including informing pathologies in tissues that are found to be aging faster, or analyzing how people with similar ages can have vastly different tissue ages. An extended technical presentation of this work can be found here, and a highly simplified non-technical overview presentation can be found here.

## 2. Background

Aging is characterized as a progressive decline in fitness that ultimately results in death. As people age, this deterioration increases the risk for many debilitating diseases, including cancer, cardiovascular disorders, and neurodegenerative conditions^1^; however, the specific mechanisms that cause aging are still unknown. ^2,3^ Thus far, one of the best known biomarkers for aging is an epigenetic clock based on age related changes in DNA methylation ^4^. While epigenetic clocks have strong predictive power, they are less interpretable since we cannot directly infer the mechanisms of aging from their predictions ^5,6^.

Therefore, we propose building tissue-specific and multi-tissue transcriptomic clocks to make global tissue age predictions. The problem of identifying tissues that age at different rates is of specific interest ^7 8^; an example application could inform pathologies in tissues that are found to be aging faster ^9^, allowing physicians to concentrate on the most vulnerable parts of the body. As a corollary, we can use the same approach to analyze deconvolved cell-specific expression data to compare aging rates between different cell types in the same tissue.

So far, classical methods such as Lasso or ElasticNet have been used on bulk expression data^10^, but we intend to improve on that by using a simple multi-layer perceptron (MLP) clock on deconvolved GTEx data. An MLP is a feed-forward artificial neural network which uses multiple layers of perceptrons, i.e., individual classifiers used in supervised learning. MLPs are particularly promising in their demonstrated strong predictive power, as they can encode non-linear relationships between the input and output data, such as the interaction of multiple genes impacting the age.

We also used XGBoost, a boosted random forest classifier. This model, released in 2016, uses a collection of decision trees that divide on subsets of features, where bootstrap aggregation is used to aggregate the final outputs of each decision tree into a final answer. This model is popular, as it provides interpretability and feature importance, while outperforming many traditional models ^13^.

Past methods have primarily analyzed only single tissue models ^11^, making it a poor predictor of different tissues with very different cell composition. Note that there is an analog between a predictor trained on non-deconvolved data across multiple tissues to predict human age, and a predictor trained on deconvolved data across multiple cells within a tissue to predict tissue age. Namely, because the data is in the same format (a matrix of RNA expression levels for tissues or cells), we can use the same model architecture trained on these two different modalities of data.

Thus, we attempt to train on pairs of tissues and all the tissues, but we accept the tradeoff that fewer patients have multiple tissues. We also used a feature importance map to identify genes that are most strongly correlated with predicting tissue age. In the future, these newly-identified genes can be used to compare aging between different tissue types and determine how rapidly tissues age relative to each other, an important source of information for clinical diagnosis revisited significantly in the last 20 years ^2,12^.

## 3. Aims

### 3.1 Aim 1

Build transcriptomic clocks which can predict the age of a person given gene expression in various tissues with an MLP. Our goal is to improve existing prediction performance, and we will also try to use deeper models such as MLPs and XGBoost to make better predictions than a simple linear regression, as it has been shown to be effective ^7,11^. We will build both tissue-specific and multi-tissue clocks for this purpose.

### 3.2 Aim 2

From our models, extract the most significant genes contributing to age prediction. We can then validate our findings against known genes linked to aging to directly infer associated mechanisms.

### 3.3 Aim 3

Analogously, explore the building of transcriptomic clocks which predict the age of a tissue given gene expression data in its individual cell types. We plan to use newly deconvolved GTEx data, and aim to improve upon the performance of the existing model of a tissue specific ElasticNet with bulk expression data.

## 4. Methods

### 4.1 Dataset Pre-Processing

#### 4.1.1 GTEx Dataset

Our data is primarily derived from the GTEx dataset (https://gtexportal.org/home/), a matrix of genotype-tissue expression values from 54 different tissues. Tissues in GTEx range from a few dozen samples to thousands of samples. Each tissue is labeled with specific variables regarding the donor, including age, sex, ethnicity, RNA integrity number, cause of death, and time after death. Gene expression is shown in transcripts per million (TPM), calculated as

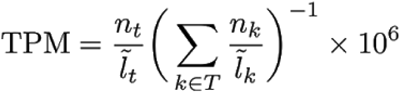

where n_t_ is the number of reads for a gene t, l_t_ is the normalized transcript/gene length, and T is the set of all genes depending on whether the quantification is at the gene level.

We directly performed inference on the expression values in GTEx, and used the chronological age of each person given as a ground truth for the tissue-specific ages as well. Despite some serious pathology notes for particular individuals, the GTEx dataset samples tissues from a population normal relative to others of their age, so these ages can be used to approximate the ages of tissues ^5,14^.

#### 4.1.2 Deconvolved Data

In addition to this bulk data, Kai Kang from the Kellis Lab provided us with computationally deconvolved data, by separating tissue expression values by individual cell types. Kang’s work used implemented deconvolution with a novel complete deconvolution method called CDSeq. CDSeq is a reference-free method that can distinguish which reads come from which cell types without needing prior experiments. This differs from reference-based methods like CIBERSORT by eliminating bias towards how the cell type was identified by previous researchers.^15^

CDSeq uses bulk RNA-seq data where the profile represents a weighted average of the expression profiles of the constituent cell types. It then takes the bulk RNA-seq data for a collection of samples and performs complete deconvolution that simultaneously outputs the cell-type proportions for each sample and cell-type-specific expression profiles. This deconvolution method extends the original latent Dirichlet allocation model in two key ways. First, the random variable that models cell-type-specific expression profiles depends on the gene length. Second, the probability of producing a read from a cell type depends on the proportion of that cell type in the sample and the typical amount of RNA produced by cells of that type. This accounts for the possibility that different cell types produce different amounts of RNA and therefore eliminates bias in the cell-type proportion estimates.^16^

### 4.2 Reproduce Existing Results, and Integrate New Data

Our code is built off of code from Elvira Kinzina in the Kellis lab. We extend her Python code for the linear regression model currently used to predict age from GTEx data. We compared our new models to the performance of her original model. To reproduce Elvira’s existing results and set up her existing pipeline for integration of our new methods, we coded in Python 3.6 in Jupyter notebooks, downloading all package dependencies which can be found in our attached code. Code can be accessed at Github here^1^.

Note that original data in the .pkl format and new data in the .RData file format required standardization to allow for direct analysis in the same pipeline.

### 4.3 Develop Tissue-Specific Predictors of Age Using Bulk Expression Data

Data was formatted as a matrix of tissues by genes for each person, expressing the level of RNA-Seq gene expression in each person’s tissue. We then ran various machine learning methods such as random forests with XGBoost and MLPs on this data. XGBoost allowed greater interpretability by determining the set of genes that appear in the most decision tree nodes. Once we had a model trained on tissues such as adipose, heart, brain, etc., we ran a grid search hyperparameter tuning with RandomizedSearchCV from scikit-learn. Tuned parameters include number of trees and depth of trees, and optimal values can be found in the code.

We also attempted an MLP trained on each patient’s genes. However, the models did in general comparably or worse to XGBoost, and varied widely from run to run, possibly as a result of not being able to run to convergence on a full hyperparameter grid.

### 4.4 Develop Multi-Tissue Predictors of Age Using Bulk Expression Data

We trained MLPs and XGBoost models by building multi-tissue transcriptomic clocks, to try to improve upon tissue-specific model performance. We first built a model using all 8 tissues of focus in the current study. However, as further discussed in the results section, only 93 individuals in our dataset had samples taken for all these tissues. The lack of training data for patients with multiple tissues resulted in poor predictive power of the multi-tissue clocks.

To help mitigate the issue of small datasets, we also built multi-tissue predictors with pairwise tissue data combinations, and compared them to the single-tissue clocks.

### 4.5 Extract Most Important Features/Genes for Age Prediction

For ease of analysis, we used XGBoost’s built-in plot_importance function on our tissue-specific clocks to extract the genes/features the models believed to be most relevant to predicting age. This function calculates importance by using the average gain of splits in trees which use the feature. Further discussion of our choice of this metric is found in section 5.2. We then performed a literature review to validate some of the genes which have been identified against genes known to correlate with age.

### 4.6 Explore Use of Deconvolved Cell Type-Specific Data for Prediction

Deconvolved GTEx data was given to us in the format of a 3D matrix for each sub-tissue: [imputed gene expression values] x [patient sample IDs] x [cell types]. The values of the matrix were the gene expression count (in TPM) for a selected gene in a selected cell type. To transform the deconvolved data into a usable format for the MLP, we transformed the 3D matrix into a 2D matrix of genes x cell types matrix for each sub-tissue sample. The deconvolved data also included a matrix with proportions of each cell type per sample; this could be directly inputted without manipulating the data format.

Since deconvolution is a computation-intensive process, we were only able to obtain deconvolved data from Kang et al. for a small number of samples for very few tissues, namely subtissues only for the brain and heart. More specifically, the brain data only had around ∼150 samples per sub-tissue (amygdala, hippocampus, etc.) and the heart data only had ∼400 samples per sub-tissue. As such, we could only perform some preliminary analysis, but there remains potential for further future investigations.

## 5. Results and Discussion

### 5.1 Aim 1: Building Tissue-Specific and Multi-Tissue Transcriptomic Clocks

#### 5.1.1 Tissue-Specific Clocks

To evaluate the performance of our models, we first benchmarked against an existing baseline. Building off of the work done by Elvira, the blue bars in Figure 1 show the performance of her existing model, operating as a combination of PCA and other preprocessing, along with LASSO and ending with linear regression. To achieve better training results, we only analyzed the tissues in the GTEx dataset with the most samples collected, which are, in order: brain, skin, esophagus, blood vessel, adipose tissue, blood, heart, and muscle. As we are comparing the age predictions made by the model to the actual values, we used the Pearson correlation coefficient *r* as an evaluation metric for our models.

**Figure 1.**
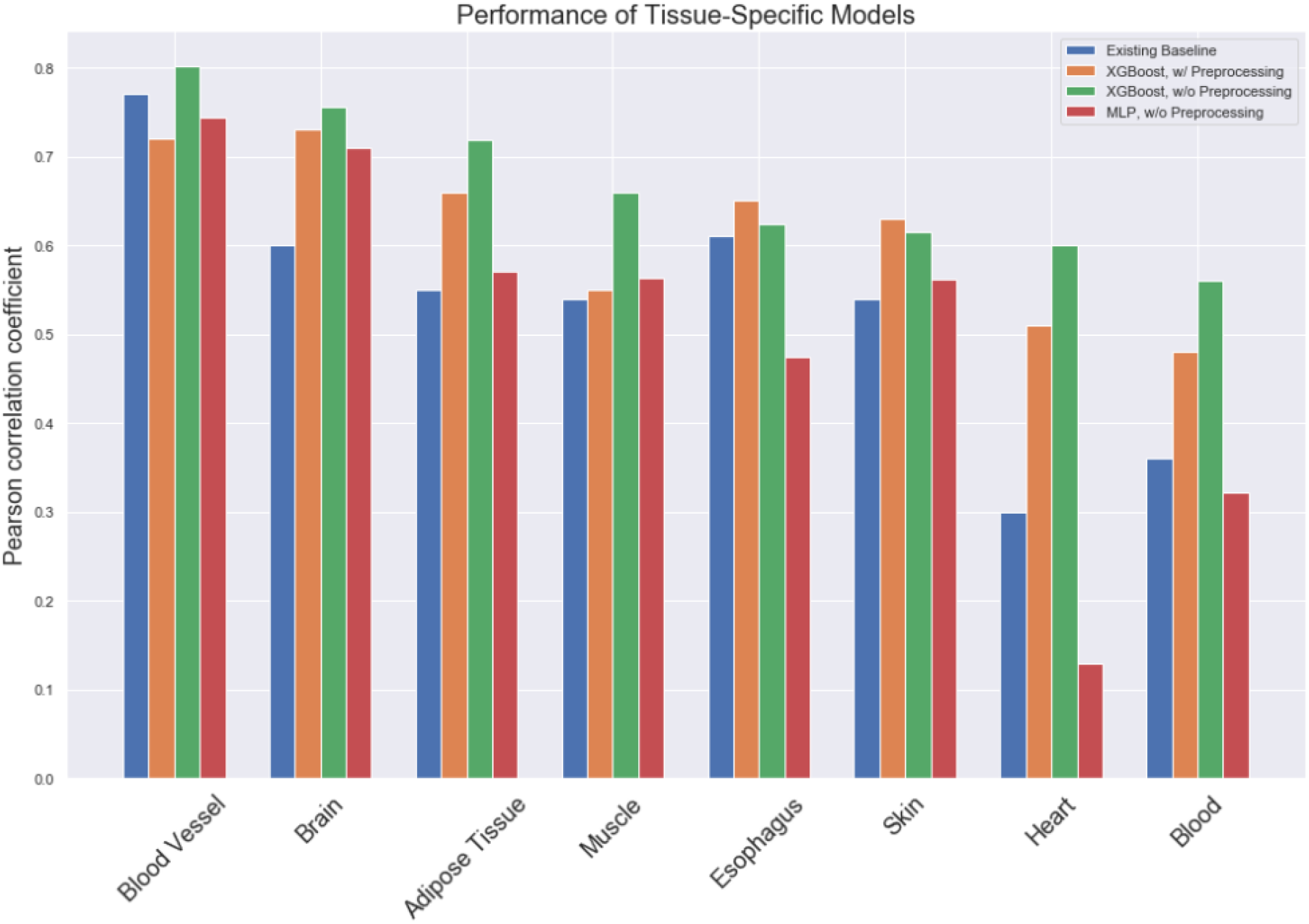
Comparison of performance of various tissue-specific models, measured by Pearson coefficient.

We then built our XGBoost model on these same tissues, first performing the same preprocessing as Elvira before, including PCA and outlier removal. With this processed data, we achieved the results in Figure 1, in orange. We note that we observe significant improvement in all tissues compared to above, except in blood vessels.

Now, to seek further improvements and interpretability, we then directly fed the raw GTEx expression data into a new XGBoost model, without the additional preprocessing as done previously. We hypothesized that this additional data would allow for stronger predictive power, and we could directly use the features themselves to infer genes and thus mechanisms correlated with aging. Our resulting Pearson correlation coefficients are found in Figure 1, in green; we note significant improvements in performance in almost every tissue compared to the above two models. Some sample plots showing the relation between predicted and actual age values are shown below in Figure 2.1 for our best tissue-specific XGBoost models.

**Figure 2.1.**
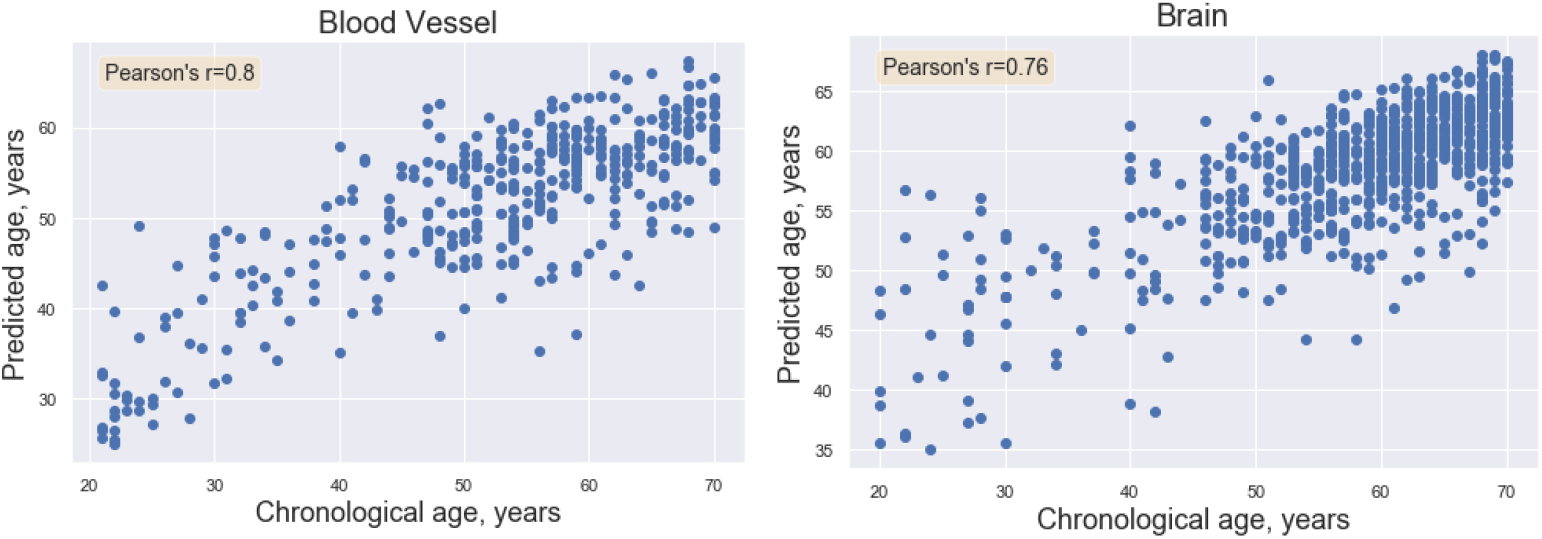
Sample plots showing the relation between predicted and actual age values for XGBoost clocks.

We also experimented with using an alternate MLP instead for the tissue-specific clocks; these results are shown in Figure 1, in red, and sample predictions are below in Figure 2.2. The structure found by the hyperparameter search on the blood tissue is two hidden layers with size of 100 and 10, with adaptive learning rate 0.0003 and regularization alpha of 0.0001, for about 3000 epochs. We hypothesized that it would do slightly worse than XGBoost, and that was indeed the case -- while it was often comparable, it was occasionally much worse in addition to being less interpretable. The performance of this model on the muscle and blood vessel data is shown below. Due to our experience with MLPs, we note that this relatively poor performance is most likely due to the very low amounts of data available to us for training, as MLPs achieve greater performance with increased training data. Given our data and performance limitations, we conclude that we will continue to next steps with only the XGBoost model framework.

**Figure 2.2.**
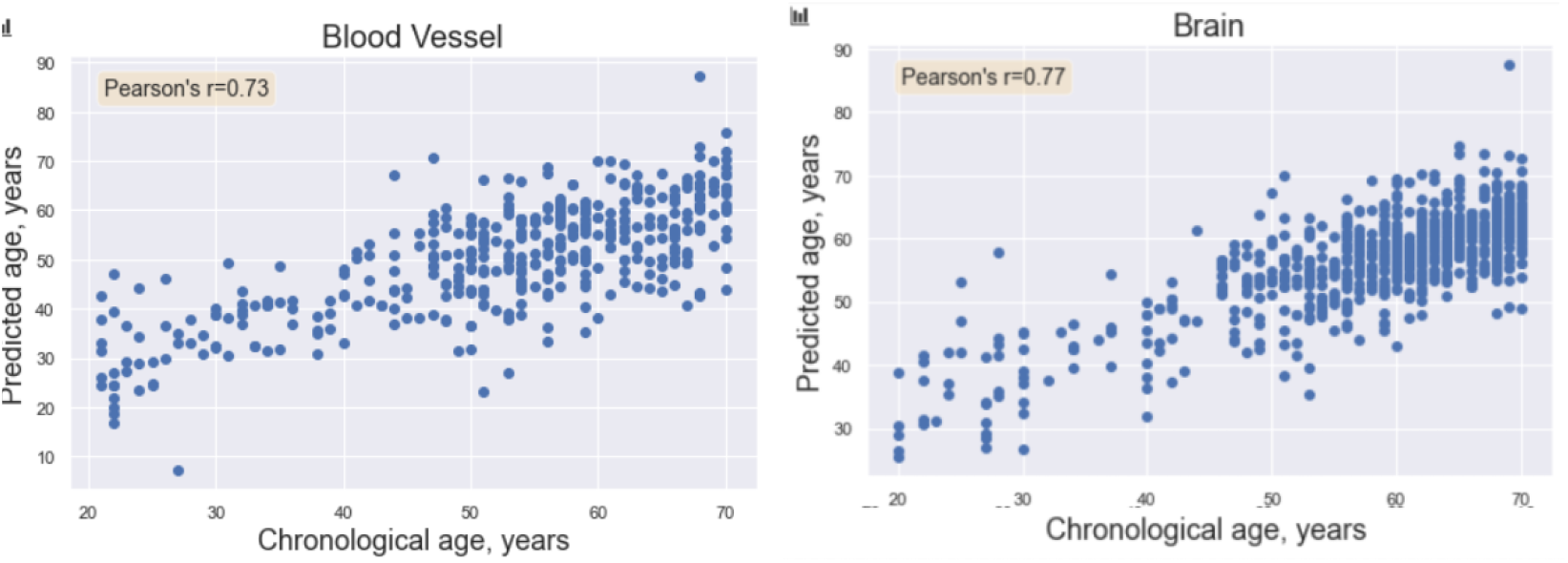
Sample plots showing the relation between predicted and actual age values for MLP clocks.

#### 5.1.2 Multi-Tissue Clocks

To build multi-tissue clocks, we used XGBoost and MLPs on all tissues, and pairs of tissues. Using XGBoost, we first experimented with building multi-tissue clocks with individuals having expression data across all 8 tissues explored above. However, there were only 93 individuals in our GTEx dataset where this is the case, so we did not anticipate strong results. The performance of our resulting, tuned model is shown in the scatter plot in Figure 3. With a correlation of 0.56 in our sparse dataset, we note that the performance is approximately equal to that of the weakest tissue-specific model above, i.e., the blood-specific clock.

**Figure 3.**
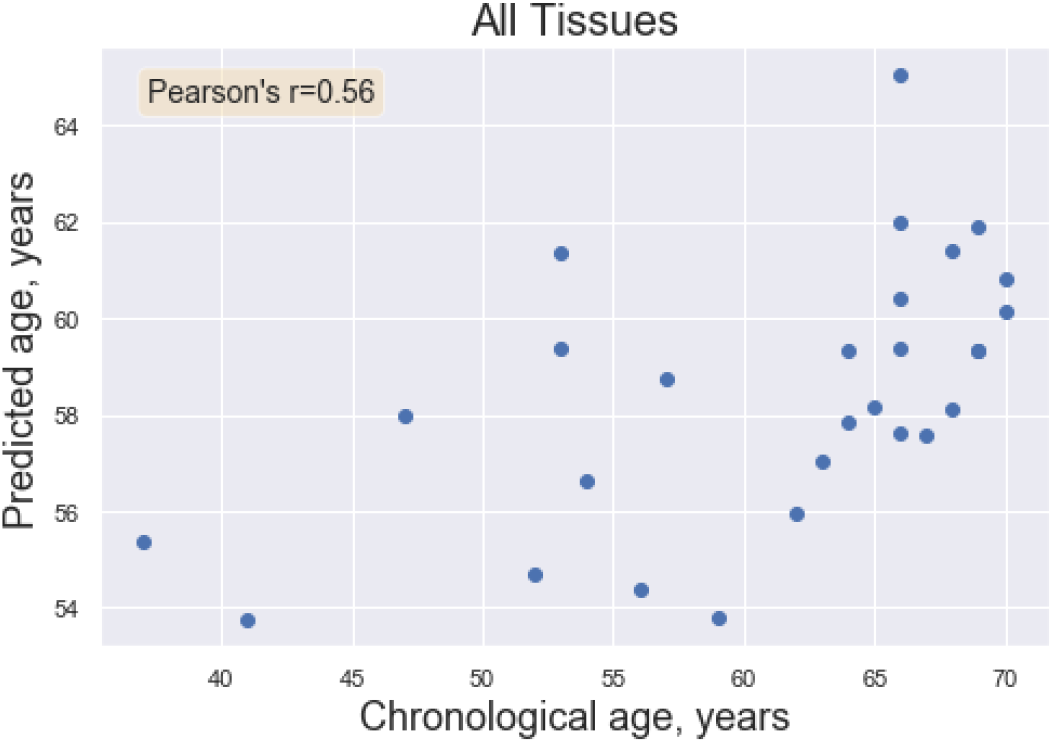
Plot demonstrating the performance of the all-tissue XGBoost model.

While we did not see improved performance as compared to the best tissue-specific model above, we note that this performance was achieved on an incredibly small sample size relative to those typically used in machine learning. This seems to suggest that this multi-tissue approach holds promise, but will require larger and more extensive datasets to better understand if this approach can perform more strongly.

To explore a compromise between entirely tissue-specific and entirely multi-tissue clocks, we explored the performance of these models when tissues are combined pairwise, and compared this performance to predictions made using the tissues individually. The results of this investigation are below, in Figure 4.1; we took the correlation coefficient from the joint model and subtract the maximum of the correlation coefficients of the two independent corresponding tissue-specific models for XGBoost. Figure 4.2 is similar, but for the MLPs.

**Figure 4.1.**
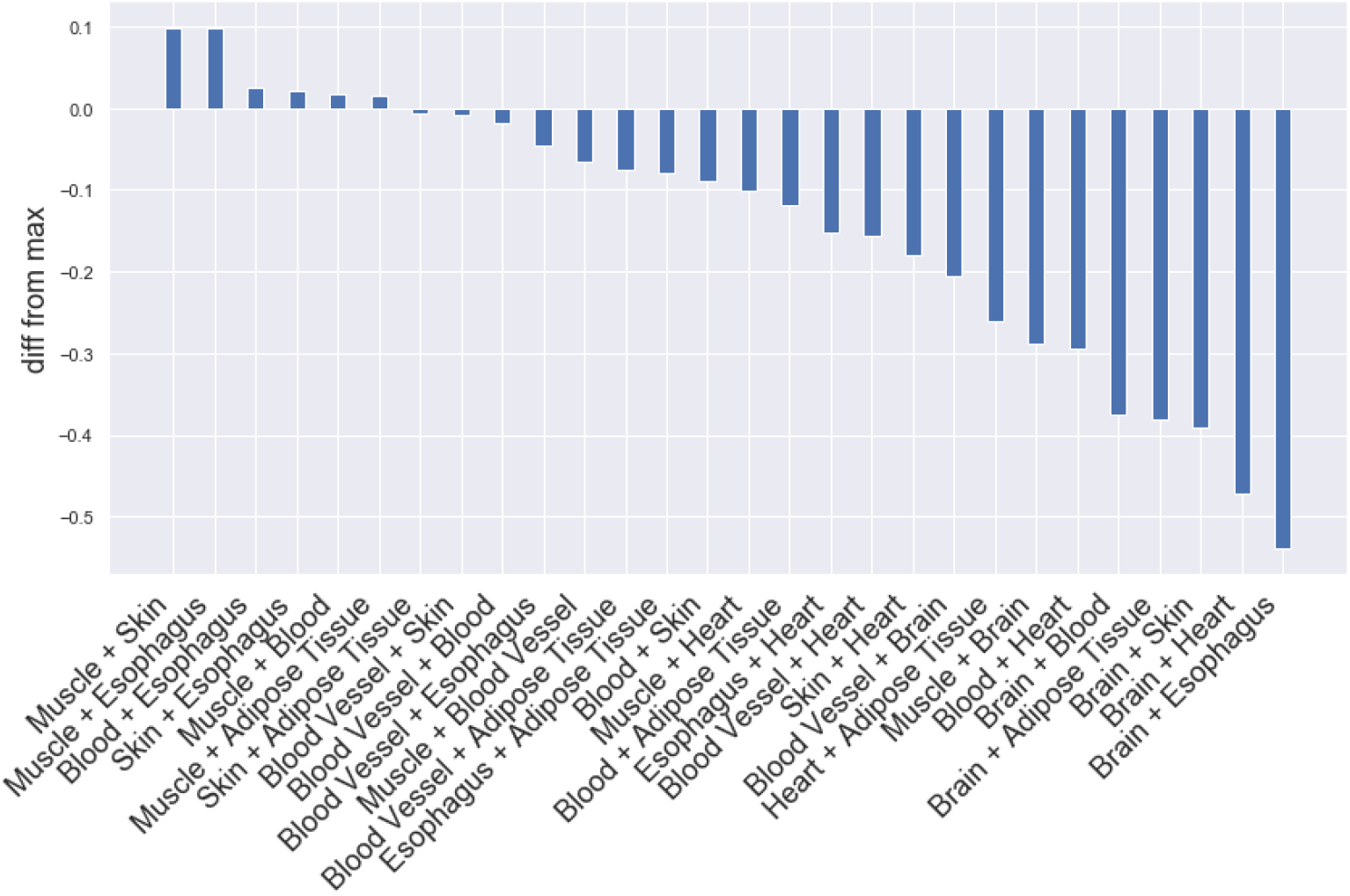
Plot demonstrating the performance of pairwise-tissue clocks in XGBoost.

**Figure 4.2.**
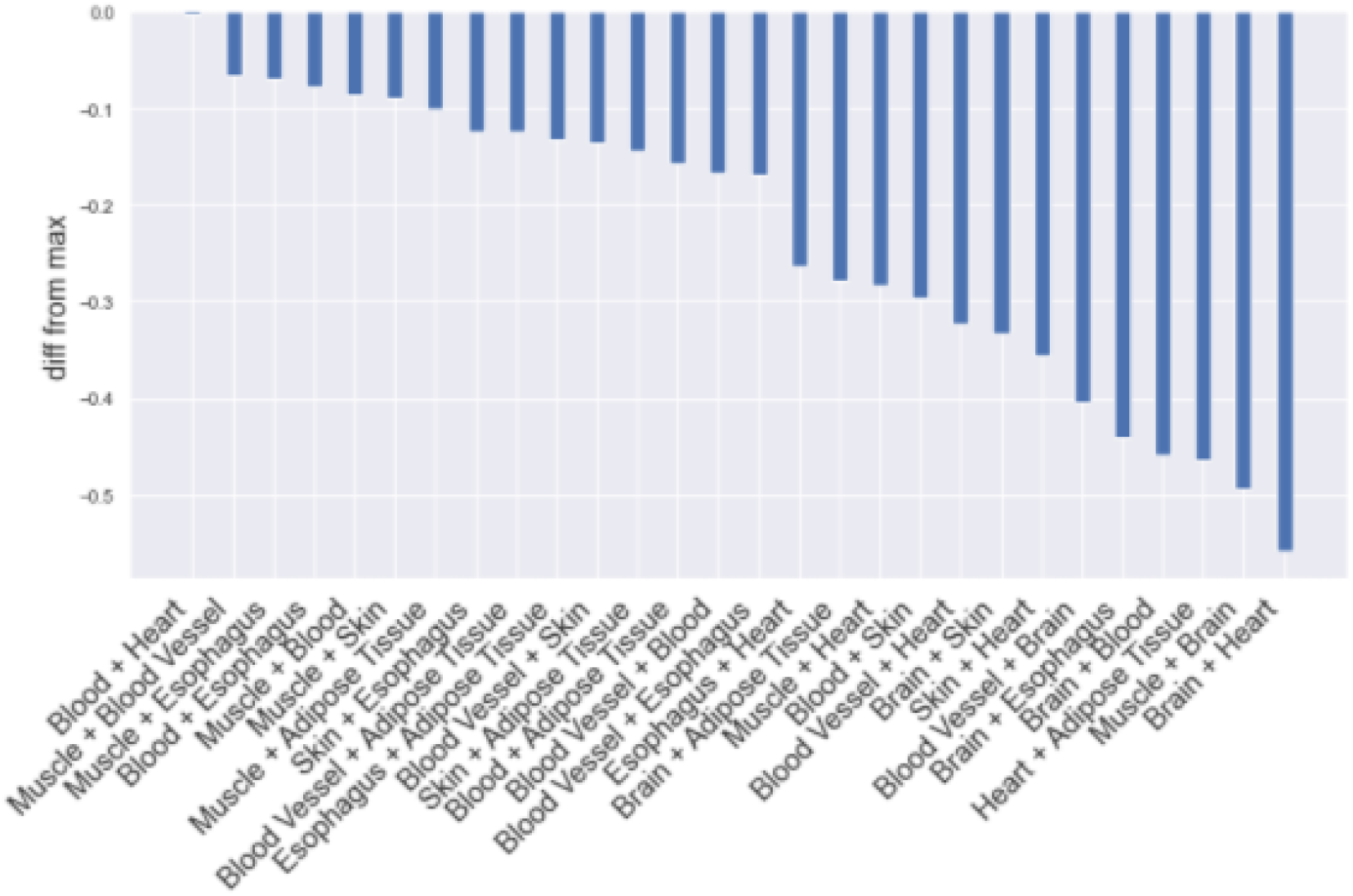
Plot demonstrating the performance of pairwise-tissue clocks using MLPs. The y-axis demonstrates the improvement upon the best single tissue clock. Note all values are negative.

These figures show that most of the models perform worse than before; MLPs do worse on every combination, and XGBoost only shows noticeable improvement on the muscle-skin and muscle-esophagus clocks. Given our limited-size datasets, it is difficult to make definitive conclusions about the multi-tissue approach; we believe the primary driver for this lack of performance is the dearth of training data across multiple tissues. However, this approach at least shows some promise and is worthy of further investigation with larger datasets in the future.

### 5.2 Aim 2: Inferring Genes Correlated with Aging

To infer the genes most correlated with aging, we exclusively used our tissue-specific XGBoost models. As XGBoost is a modified random-forest technique, the model lends itself to direct interpretability; in contrast, the problem of interpretability in MLPs still remains a relatively open question ^17,18^.

The XGBoost framework offers three methods for calculating the importance of individual features: by weight, cover, or gain^17^. Namely, weight uses the number of times a feature appears in trees, cover uses the average coverage of tree splits using the feature, while gain is the average gain of splits in trees which use the feature. Our experimentation found that the “weight” method was unreliable, in that it often simply reported genes appearing earlier in the dataset due to the nature of the XGBoost algorithm, and the “cover” method gave almost the exact same results for all the top genes in the model. As such, we used the “gain” method, which we found to be most interpretable and also the most intuitive as a measure for how important the genes are to the prediction. If using a particular gene in making predictions gives better performance, that gene seems likely to be an important marker of aging.

With our tissue-specific models, we then analyzed the models to extract the genes most important to the models for predicting age; the top four genes for each tissue can be found in Table 1 above. Across all eight tissues, there was one common gene between the muscle and heart: EDA2R. EDA2R is a gene that encodes a transmembrane protein in the tumor necrosis factor receptor superfamily^19^. Previous research that investigated disease gene enrichment for core-aging genes (CAGs), health-specific aging genes (HSAGs), and common-specific aging genes (CSAGs) found that EDA2R was one of the top up-regulated CAGs in the tibial artery tissue and was therefore considered a strong candidate gene for lung aging ^19,20^. EDA2R has also been associated with chronological age using SOMAscan, a process that uses SOMAmers (single-stranded DNA aptamers) to quantify the level of subproteome of proteins ^19–21^.

**Table 1.** Table showing the top four genes for each tissue in order from left to right. Gene highlighted in green, EDA2R, was found to be important in both the muscle and heart tissue-specific models. Genes highlighted in yellow (GABRB1, SUMO4, HGH1) have known links to aging.

| Tissue | Most important genes (in order from left to right) |  |  |  |
| --- | --- | --- | --- | --- |
| Muscle | EDA2R | GALNTL6 | CXorf57 | LANCL1-AS1 |
| Blood Vessel | RP11-731J8.2/GABRB1 | PTCHD4 | TSEN34 | HCG11 |
| Brain | KIF3B | HPCAL4 | SMIM17 | GRIK1 |
| Blood | LINC00957 | RP11-960L18.1 | GIPR | TRUB2 |
| Skin | NAT8L | SNAPC1 | LHFPL4 | GDF15 |
| Esophagus | SUMO4 | AC068831.6 | RP11-373D23.2/FOSL2 | ZNF649 |
| Heart | HGH1 | ARMCX6 | STARD5 | EDA2R |
| Adipose Tissue | PYHIN1 | MTHFD1L | PCK1 | CXCR4 |

Other extracted genes with links to aging include GABRB1, SUMO4, and HGH1 (Table 1). First, GABRB1 is a component of GABA, the major inhibitory neurotransmitter in the brain. It is the target of the drug Zolpidem (more commonly known as Ambien); since sleep and memory both decrease with age, a Phase IV trial is currently studying Zolpidem to see if the drug can increase memory-related sleep events and improve post-sleep memory recall in older adults ^19–22^.

Second, SUMO4 encodes a small, ubiquitin-related modifier that controls the target proteins’ subcellular localization, stability and activity. Researchers have previously found that aging alters SUMO4 expression in adipose-derived stem cells, and that aging causes a significant increase of SUMO levels in *C*.*elegans* ^*23*^. SUMOylation, or the modification of proteins by the ubiquitin-like protein, has also been found to directly influence aging and senescence, the irreversible loss of cell replication that occurs during aging ^23,24^. As individuals age, there are higher levels of global protein SUMOylation and a decrease in the components of SUMOylation process ^23–25^.

Finally, HGH1 is a human growth hormone that is produced by the pituitary gland and stimulates physical growth and cell regeneration. Growth hormones naturally decline during aging, inducing somatopause and causing decreased lean muscle mass, increased adiposity, and lower energy. However, the decrease in growth hormone levels could also protect the body from cancer and other age-related diseases and increase patient lifespan ^23–26^. Overall, our models have identified genes that are known and validated to play key roles in tissue-specific aging, disease risk, and physiological fitness. As such, it is likely that other genes identified by our models, which have not previously been studied for linkage to aging, can yield promising candidates for further investigation.

We note that the fact that we had one matching gene among the top 4 genes reported for each tissue is a very promising result. We recognize that the most important genes in each tissue should generally be different, due to their heterogeneity and differentiation in function and gene expression, but this match appears to serve as validation of our results. Despite differentiation in expression, it is expected that some important genes are still generally shared between tissues, and given that the heart is also a muscle, the shared EDA2R gene between the two tissues is noteworthy.

To determine the significance of these results, we calculate the probability of having no matches among the top 4 genes if they were picked by random chance from each tissue. It can be computed as follows: given 20000 genes per tissue, the probability of selecting 32 unique genes between 8 tissues is equal to the below expression, approximately equal to 0.978:

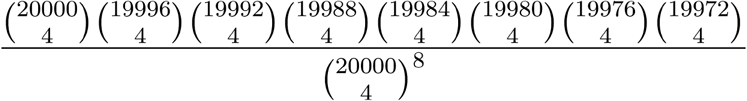

This gives only a 0.022 chance that there is at least one match by random chance, so this suggests that our models are capturing statistically significant biological information.

### 5.3 Aim 3: Exploration of Use of Deconvolved Data

Our final results were computed on the deconvolved cell-specific data from Kang et. al; these results are shown in the plot below in Figure 6. The deconvolved data was only available for these particular subtissues due to computational time constraints.

**Figure 6.**
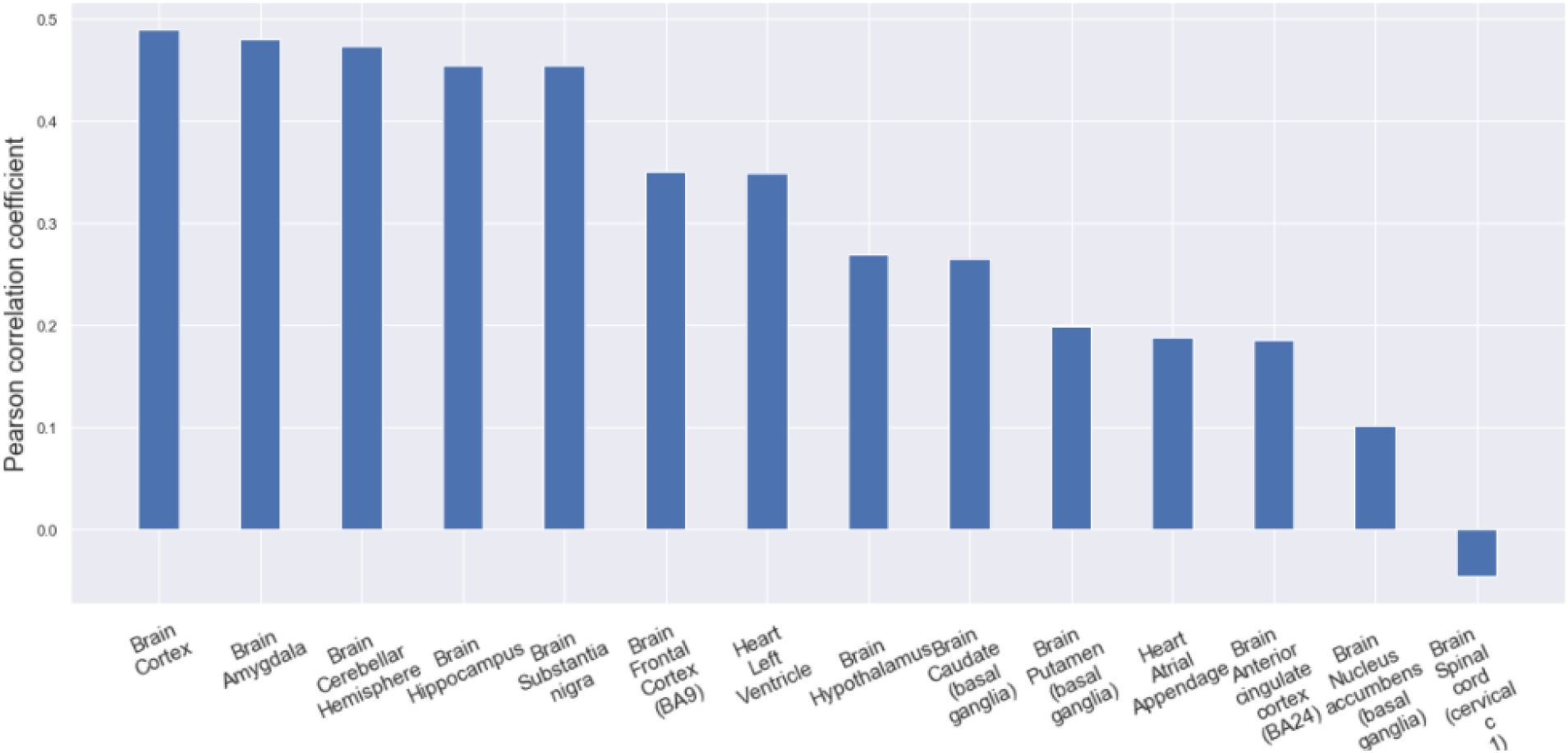
Plot demonstrating the performance of XGBoost on deconvolved cell data for various subtissues.

We noted that the best models here have correlation coefficients less than 0.5, which is worse than the corresponding brain and heart models in our tissue-specific models earlier in our investigation. However, the amount of data used in our largest deconvolved data models was only 20% of the size of our smallest whole-tissue datasets. We also noted that there did seem to be particular subtissues within the brain and heart which served as better predictors than others, suggesting that this technique’s power may be somewhat tissue-specific.

As such, having performance which appeared to approach that of our earlier whole-tissue models with significantly less data seems to suggest that this method could potentially hold promise for future investigation, given sufficient data. However, our investigation was unable to yield any additional improvements in our model performance beyond our existing whole-tissue models, so we did not observe deconvolution leading to reduced noise and better predictions for the time being. This is likely because deconvolution leads to a significant magnification in the amount of data the models needed to analyze to find meaningful relationships, suggesting a tradeoff: deconvolution can generate more data to reduce the effects of heterogeneity of cell-specific expression, but the increase in data also makes it more challenging for models to perform well on such limited datasets.

## 6. Future Work

We currently predict age for each tissue in each individual with our model. As a result, the clocks are slightly different per person (ex: some people have an older-looking liver, others have a younger-looking brain) and produce an aging rate different than what we’d expect if all tissues were aging at the same rate. Kellis mentioned that people with diseases in specific tissues (such as the liver or brain) might show traits like this, and we’d like to investigate his hypothesis further. Researchers have previously used the GTEx dataset to create a “healthy” aging cohort that included donors who did not die due to tissue-specific, chronic diseases. Age-associated genes derived from this cohort were compared with those from the “common” aging cohort and found that both cohorts shared a large number of aging signatures. However, “common” aging genes were found to be driven by the regulation of age-related disease genes, while “healthy” aging genes were more strongly regulated by protective mechanisms that prevented disease development. Disease gene enrichment for the “healthy” and “common” aging signatures also varied in a tissue-specific manner ^20,23–26^. Therefore, incorporating phenotypic information for GTEx into the models could lead to findings that individual pathology can be indicative of tissue-level aging.

Kellis also suggested that telomere length could be correlated with tissue age. Telomeres shorten with age and telomere shortening can lead to senescence, apoptosis, genomic instability, and oncogenesis. Older people with shorter telomeres have a three-times higher risk of dying from heart disease and eight-times higher risk of dying from infectious diseases. The rate of telomere shortening therefore directly impacts an individual’s health and pace of aging. Furthermore, habits like smoking, limited physical activity, high stress, and unhealthy diet can increase the rate of telomere shortening ^27^.

If telomere length is available for a subset of individuals in a subset of tissue, we could use the telomere length-based age predictions as the “gold standard” for training or validating our clock’s predictions in each tissue. This could involve directly training a model with telomere length to predict age in each tissue independently, then comparing the results to a model trained on chronological age. We could also train a tissue-specific age prediction model from telomere length, and then use the telomere-length based prediction as the age for the expression-based model. In other words, we could use a variant of transfer learning from telomere length predictions to improve predictions in our gene expression model.

## 7. Acknowledgements

We would like to acknowledge Manolis Kellis, as well as Elvira Kinzina and Kai Kang from his lab at MIT.

They provided valuable feedback on ideas, project direction mentorship, and dataset access. We also would like to acknowledge Lei Hou and Xushen Xiong in the Kellis Lab for helping us to better understand various aspects of the GTEx dataset.

## Footnotes

1 https://github.com/mdlu/aging-clock

